# Spatial modeling of the membrane-cytosolic interface in protein kinase signal transduction

**DOI:** 10.1101/191940

**Authors:** Wolfgang Giese, Gregor Milicic, Andreas Schröder, Edda Klipp

## Abstract

The spatial architecture of signaling pathways and the interaction with cell size and morphology are complex but little understood. With the advances of single cell imaging and single cell biology it becomes crucial to understand intracellular processes in time and space. Activation of cell surface receptors often triggers a signaling cascade including the activation of membrane-attached and cytosolic signaling components, which eventually transmit the signal to the cell nucleus. Signaling proteins can form steep gradients in the cytosol, which cause strong cell size dependence. We show that the kinetics at the membrane-cytosolic interface and the ratio of cell membrane area to the enclosed cytosolic volume change the behavior of signaling cascades significantly. We present a mathematical analysis of signal transduction in time and space by providing analytical solutions for different spatial arrangements of linear signaling cascades. These investigations are complemented by numerical simulations of non-linear cascades and asymmetric cell shapes.

## 1 INTRODUCTION

Cells need to respond to a large variety of external stimuli such as environmental changes or extracellular communication signals. Signals transmitted from cell surface receptors to target genes in the nucleus are frequently transduced by cascades of covalent protein modifications. These modifications consist of inter-convertible protein forms, for instance, a phosphorylated and an unphosphorylated protein. Signaling cascades occur in many different variations including mitogen-activated protein-kinase (MAPK) cascades and small GTPase cascades.

Signal transduction mechanisms carried out by networks of protein-protein interactions are highly modular and regulatory behavior arises from relatively simple modifications Bhattacharyya et al. (2006). The spatial arrangement of signaling cascades varies in different biological systems. We focus on the localization of signaling components, which can be tethered to the cell-membrane or freely diffuse in the cytosol. Tethering to the cell-membrane can be mediated by protein prenylation Gelb et al. (2006); Wang and Casey (2016), co-localization by membrane-bound scaffolds Gordley et al. (2016) or membrane anchoring proteins Alberts et al.. Frequently, the first steps of signal transduction occur at the membrane and are then continued into the cytosol.

In many experimental and theoretical studies on signaling cascades, the cell is regarded as a number of well-mixed compartments with no variation in size, shape or organelle location. An extensive mathematical analysis of temporal aspects of signaling processes has been carried out in quantitative biology Heinrich et al. (2002); Kofahl and Klipp (2004); Klipp and Liebermeister (2006); Beguerisse-Díaz et al. (2016). However, the spatial description of signaling processes has received less attention despite its relevance in understanding cell morphology and growth regulation in time and space Kholodenko et al. (2010). Examples of spatial effects on the length scale of single cells range from the yeast mating process Maeder et al. (2007); Dudin et al. (2016) to the propagation of spatial information in hippocampal neurons which is controlled by cell shape and *vice versa* Neves et al. (2008); Chay et al. (2016).

We investigate linear signaling cascades with different realizations of spatial arrangements of signaling components as shown in Figure 1. Here, we focus on the membrane-cytosolic interface, which is included in the signaling motif shown in Figure 1(**B, C**).

**FIGURE 1.**
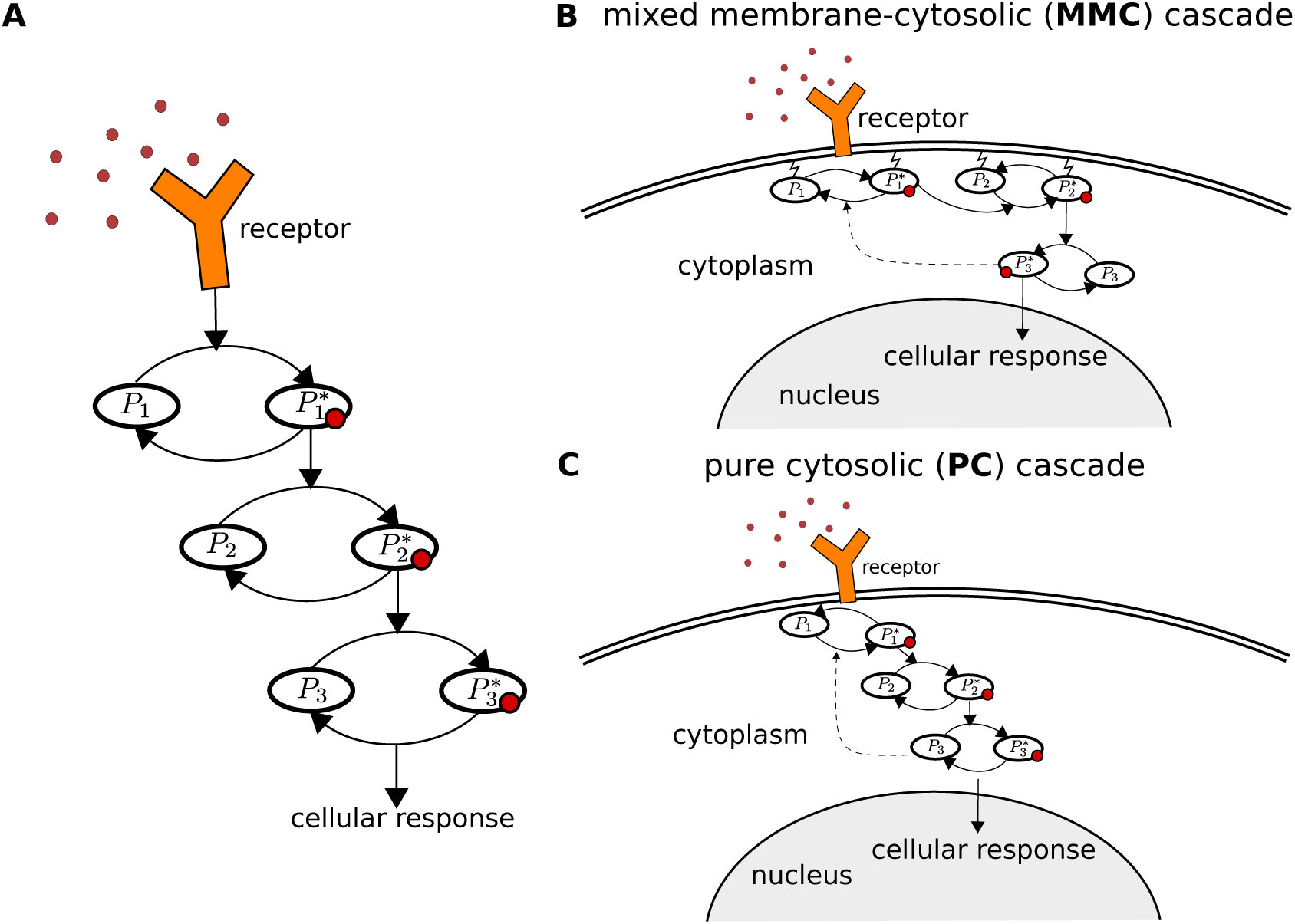
Spatial organization of signaling cascades. **A:** Sketch of the classical temporal signal transduction model. Extension of this model into three-dimensional space naturally results in a variety of different spatial motifs. **B:** The signal is first processed by signaling components tethered to the membrane, and then transduced at membrane-cytosolic interface into the cytosol. **C:** The signaling components are directly activated at the membrane-cytosolic interface and diffuse through the cytosol. Note, that diffusion coefficients for lateral diffusion along the membrane are much lower than in the cytosol.

Since the cytosol scales with cell volume and the cell membrane with the cell surface, reactions on the membrane and in the cytosol scale with the cell-surface to cell-volume ratio. For instance, we obtain an area/volume ratio of ∝ 3/*R*_cell_ for a spherical cell geometry, where *R*_cell_ is the cell radius. We will show that this affects the global phosphorylation rate of signaling proteins that diffuse in the cytoplasmic volume, which depends on cell size. While cytosolic gradients naturally occur from the membrane to the nucleus, membrane-bound components can only form gradients along the membrane, which changes the response to heterogeneous signals. Furthermore, the diffusion on the membrane is much slower for membrane-bound components than for cytosolic components Klünder et al. (2013). Both of these factors are expected to largely change signal transduction properties of the pathway.

An analysis and comparison of spatial signal transduction motifs in response to spatially homogeneous and heterogeneous signals is presented in this study. The natural extension of widespread used ordinary differential equations are bulk-surface partial differential equations Elliott and Ranner (2012); Eigel and Müller (2017). Here, *bulk* refers to the cellular compartments that are represented as a volume such as the cytoplasm or the nucleus, while *surface* refers to all cellular structures that are represented as an area such as the cellular or nuclear membrane. Since their introduction to cell signaling systems Levine and Rappel (2005), bulk-surface partial differential equations have been successfully employed in several models for cell polarization Rätz and Röger (2012); Klünder et al. (2013); Giese et al. (2015); Thalmeier et al. (2016). However, membrane-cytosolic interfaces at different stages of a signaling cascade have not yet been investigated.

We start with an analysis of two different motifs with simplified linear kinetics, which allows to develop exact analytical solutions of the steady state. Both motifs differ in their cell size dependence and we show further that their behavior can drastically differ from the assumption of well-mixed compartments. The time-scaling of signal transduction is investigated using the recently introduced method of local accumulation times Berezhkovskii et al. (2010). We continue by investigating the response and sensitivity to spatially heterogeneous signals such as signaling gradients for symmetrical and asymmetrical cell shapes. In the last section, we proceed with numerical investigations of systems with non-linear Hill kinetics as well as negative feedbacks and oscillations. A Fourier analysis in time is used to provide insight into the dependency of oscillation frequency and amplitude on cell size. Depending on the spatial motif, cell size limits for the extinction of oscillatory behavior are obtained.

## 2 THE MODEL

We start with a linear signaling cascade with different localizations of the membrane-cytosolic interface as shown in Figure 1. We employ a simple cascade model from Heinrich et al. (2002), in which stimulation of a receptor leads to the consecutive activation of several down-stream protein kinases. This model is extended into space in the following. We assume a linear cascade with *N* components, where the first *M* < *N* components are localized at the membrane while the remaining *N* - *M* components are assumed to freely diffuse in the cytosol. The equations for the membrane-bound components read

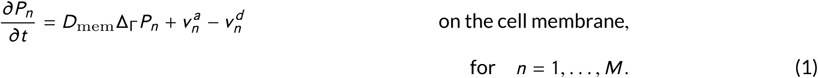

Here, 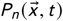 are the local concentrations on the cell membrane and 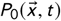 is the input signal. All of these species are functions of space and time, where 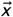 is a point on the membrane, *t* is the time and *D*_mem_ the diffusion rate on the cell membrane. Since the membrane is a surface in three-dimensional space with negligible thickness, the natural unit for concentrations of the cell membrane-bound species *P*_*n*_, *n* = 1, …, *M* is molecules per area (see Table 2). The phosphorylation rates 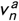 as well as the dephosphorylation rates 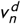 have units molecules per area and time. Note, that if the input signal is homogeneous in space, meaning 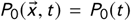, all spatial fluxes *D*_mem_ߜ_Г_*P*_*k*_ are zero and the equation system for the membrane-bound species can be described by an equivalent system of ordinary differential equations (SI text). In contrast to the membrane-bound signaling components *P*_1_, …, *P*_*M*_, the signaling component *P*_*M* +1_ can freely diffuse in the cytosol. For the modeling of the membrane-cytosolic interface, we need to include diffusion in the cytosol and reactions on its boundaries, which are the membranes. These processes are modeled by a reaction-diffusion equation

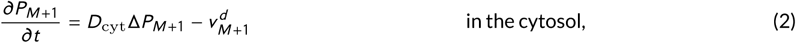

with the boundary condition

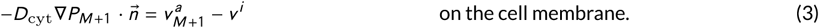

Since *P*_*M* +1_ is activated by the upstream component *P*_*M*_, which is tethered to the membrane, there is a phosphorylation reaction only on the cell membrane but not in the cytosol. This reaction is, therefore, modeled as a boundary condition. The reactions at the membrane-cytosolic interface are described by the phosphorylation rate 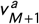 and the dephosphorylation rate *v i*, both with units molecules per area and time. The species *P*_*M* +1_ diffuses freely in the cytosolic volume with the diffusion rate *D*_cyt_ and therefore its local concentration is described in molecules per volume. The dephosphorylation rate 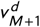 is, therefore, given in molecules per volume and time. For the flux on all other membrane enclosed organelles we assume a zero-flux condition

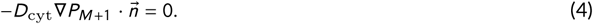

The equations for the components of the downstream cytosolic cascade read

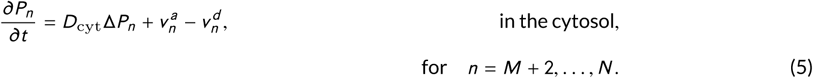

The concentrations at position 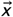 in the cytosolic volume at time *t* of the cytosolic components, 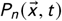 with *n* = *M* + 1, …, *N*, are given in molecules per volume (see Table 2). For the cytosolic components we assume zero-flux conditions:

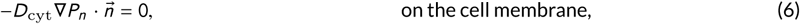

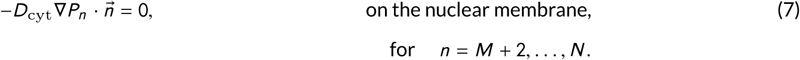

In classical MAPK cascades the last component of the cascade, which is the phosphorylated MAPK is imported into the nucleus. Examples range from Hog1 nuclear import in yeast Klipp et al. (2005); Muzzey et al. (2009) to the import of ERK in mammals Nardozzi et al. (2010). In this case the boundary condition (7) on the nucleus for the last cytosolic component *P*_*N*_ needs to be modified to

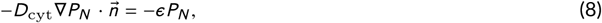

where *ε* represents a nuclear-import reaction rate on the nuclear membrane.

We will test and compare systems with three components *N* = 3 as shown in Figure 1, where the spatial arrangement of the components is varied. Here, *M* = 2 describes the case of two membrane-bound and one cytosolic element (motif Figure 1**B**) and *M* = 0 the case of only cytosolic components (motif Figure 1**C**). In the following the case *M* = 2 is referred to as mixed membrane-cytosolic (MMC) and *M* = 0 as pure cytosolic (PC) cascade.

## 3 RESULTS

### 3.1 The mixed membrane-cytosolic cascade is strongly size dependent

A spherical cell with radius *R*_cell_ is assumed in the following analysis. The nucleus is as well represented as sphere with radius *R*_nuc_, which is placed at the center of the cell. The input signal is denoted by *P*_0_(*t*) and is assumed to be homogeneous on the cell surface. The concentrations of protein kinases are described by functions *P*_*i*_ (*r*, *t*) depending on space and time. Note, since the cellular geometry is radially symmetric and the input signal *P*_0_ acts homogeneously on the cell membrane, these functions depend only on the radial distance from the origin denoted by *r* and time *t*. The model equations for the mixed membrane-cytosolic cascade (MMC, *M* = 2) with linearized kinetics read

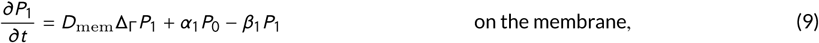

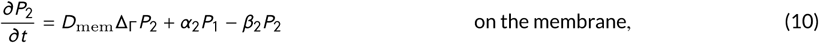

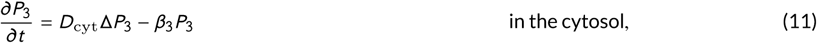

and boundary conditions for the cytosolic species *P*_3_:

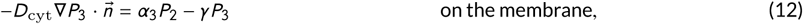

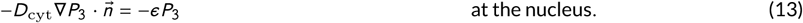

Here, *a*_*i*_ describe phosphorylation and *β*_*i*_ desphosphorylation rates. In (11), the rate *γ* with units *μm*/*s* describes saturation of the activation reaction on the membrane.

The steady state for the first two elements is given by 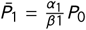 and 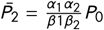. For the steady state of *P*_3_, the solution is given by

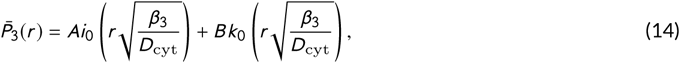

where the coefficients *A* and *B* are derived in the SI text. The steady state solution for different cell sizes is shown in Figure 2. The concentration is maximal at the cell membrane and decays towards the nucleus. An approximation of the decay length *L*_gradient_ of the intracellular gradient (with highest concentration at the membrane) is given by Brown and Kholodenko (1999)

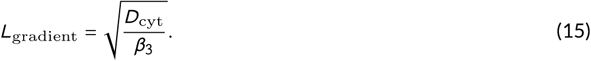

**FIGURE 2.**
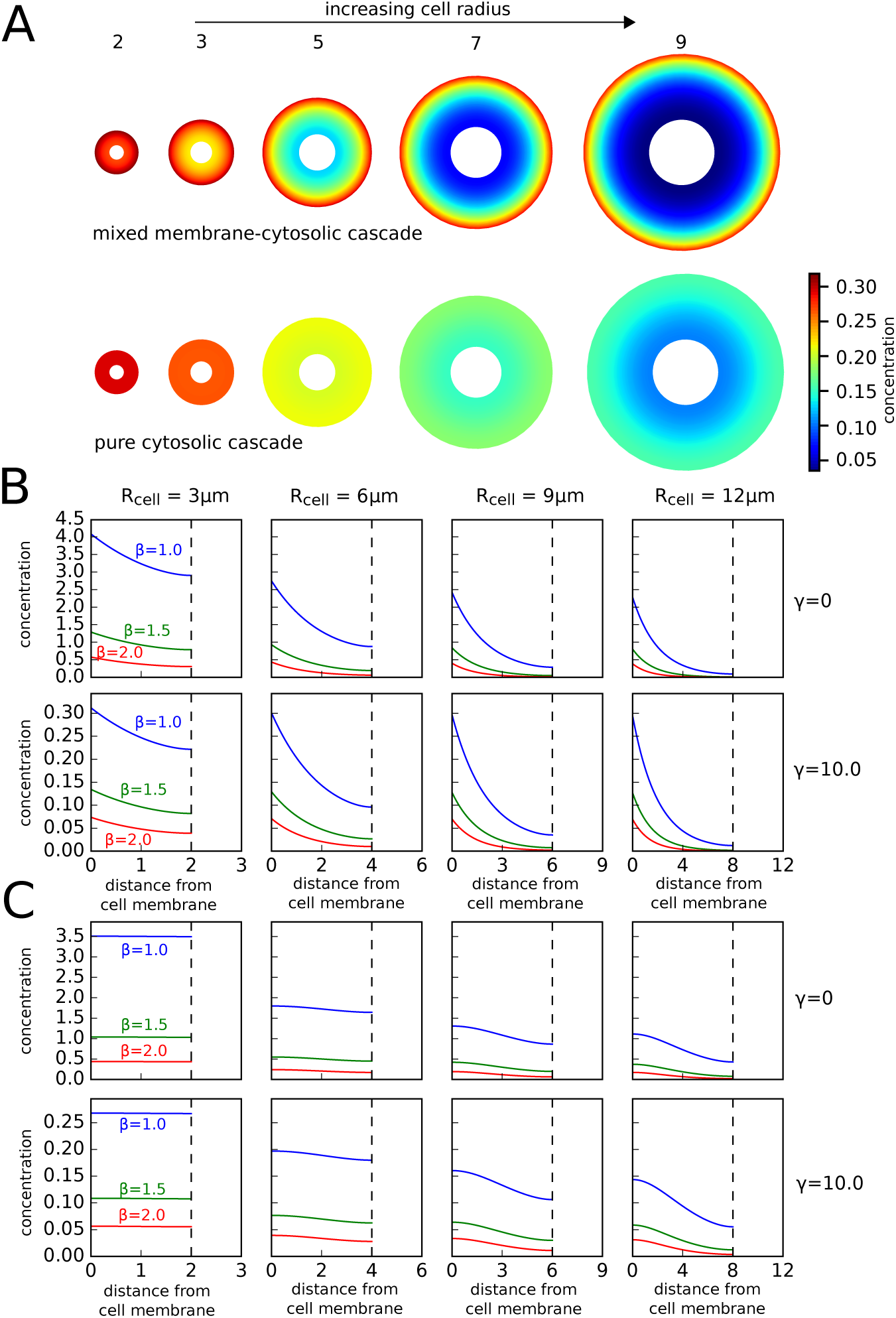
Intracellular concentration profiles for two different signal-transduction motifs. **A:** The concentration of the third cascade element *P*_3_ was plotted along a slices through three-dimensional cells. The numbers below the cells indicate the cell radius in microns, the radius of the nucleus is 1/3 of the cell radius. Strong intracellular concentration gradients are generated in the case of the MMC cascade with two membrane-bound components and only one cytosolic species [above]. Concentration gradients are much shallower for the pure cytosolic cascade, where all three components diffuse freely in the cytosol [below]. The parameters used were *D*_mem_ = 0.03, *D*_cyt_ = 3.0, *a*_1_ = *a*_2_ = *a*_3_ = 1.5, *β*_1_ = *β*_2_ = *β*_3_ = 1.0, *γ* = 10.0, *P*_0_ = 1.0. **B:** A MMC cascade with two membrane-bound components and only one cytosolic species is shown. **C:** In contrast the PC cascade, where all three components diffuse freely in the cytosol exhibits much shallower gradients. For both spatial motifs size dependence of the concentration level decreases with higher values of *γ*.

This decay length can be compared with the actual cell size. Their ratio is called the Thiele modulus, a dimensionless measure defined as 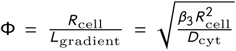 Meyers et al. (2006). For Φ > 1 strong intracellular gradients and concentration heterogeneities of signaling molecules are to be expected, while for Φ < 1 the concentration is almost homogeneous.

The effect of cell size on intracellular concentration gradients is shown in Figure 2. However, the cell size dependence in cell signaling systems does not only arise by the characteristic length scale for intracellular gradient formation *L*_gradient_ as reported in Brown and Kholodenko (1999); Meyers et al. (2006), but by the change of absolute intracellular concentration levels. The dependence of absolute concentration levels on the membrane-cytosolic interface is shown in Figure 2 and supplementary Figure S1 for a set of different parameters.

In the special case that there is no nucleus or excluding volume, *R*_nuc_ = 0, we have *B* = 0 and the steady state solution reads

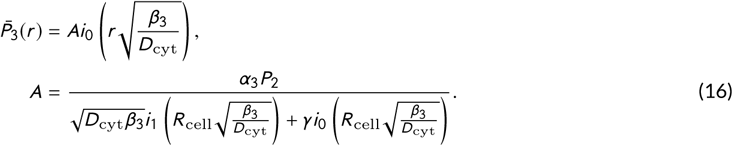

The coefficient *A* represents the minimal concentration in the cell center (*r* = 0). The maximal concentration at the cell membrane is bounded by

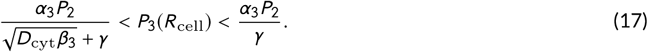

The concentration level at the membrane tends towards the upper bound in the limit of small cells, meaning Φ ≪ 1, while it tends towards the lower bound in the limit of large cells, meaning Φ ≫ 1. This effect is shown in supplementary Figure S1. Note, that for a large inactivation rate at the membrane-cytosolic interface *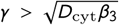*, the cell size dependence decreases. Therefore, cell size dependence is mainly determined by *γ* and *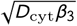*, but is independent of the phosphorylation rate *α*. In order to achieve cell size independence and a reasonable concentration of phosphorylated signaling molecules, the phosphorylation rate *α* has to be increased together with *γ*.

We can further investigate the evolution of the average concentration levels for this model in the case *γ* = 0, which implies strong cell size dependence. In case of arbitrary cell shapes with cell volume *V*_cell_ and cell membrane area *M*_cell_, the average concentration level is obtained from

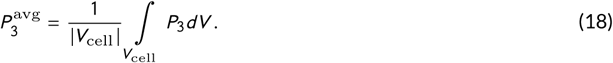

It can be shown that in this case the average concentration levels follow the system of ordinary differential equations

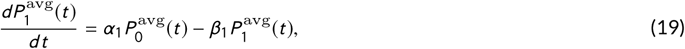

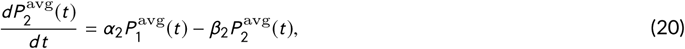

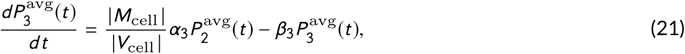

where 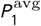 and 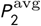 are the average concentration levels in molecules per cell membrane area. For a derivation see supplementary text S1. The steady state for the average concentration of *P*_3_ is given by

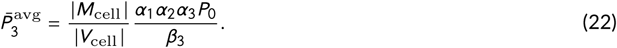

Therefore, the average concentration level scales with the ratio of 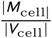. The effective global phosphorylation rate for the average concentration of active signaling molecules in the cytosol is therefore determined by 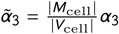. These relations give us a correspondence between widespread used ordinary differential equations and the bulk-surface partial differential equations employed in this paper. In summary, we have strong cell size dependence, with decreasing concentrations for larger cells. For cells with F Φ 1 also the concentration differences from cell membrane to cell nucleus become important and a description in terms of average concentration may become invalid.

### 3.2 Efficient cytosolic transport via cytosolic cascades

In the following we consider a PC (pure cytosolic) cascade with three elements, in which all elements freely diffuse through the cytosol. The reaction-diffusion system is given by

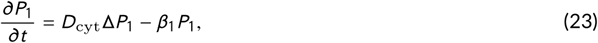

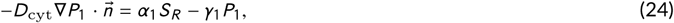

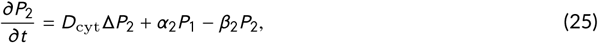

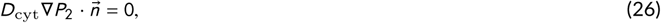

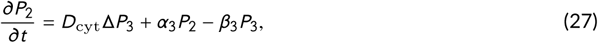

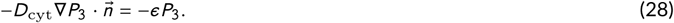

In the special case of *β*_1_ = *β*_2_ = *β*_3_ = *β* analytical approximations to cytosolic cascades in a one-dimensional system have been derived in Stelling and Kholodenko (2009); Muñoz-García et al. (2009). While a one-dimensional cellular geometry can be used to study gradient formation qualitatively, spatial effects such as the cell surface to volume ratio are neglected. Therefore, we present exact analytical solutions to the linear system in three dimensions. The steady state solutions for 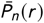 are expanded as follows

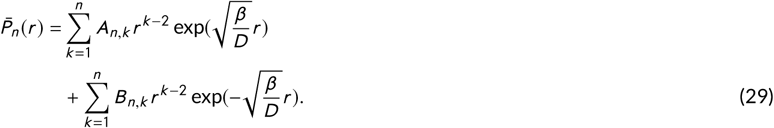

The algebraic expressions of the coefficients *A*_*n*,*k*_ and *B*_*n*,*k*_ and their derivation are shown in the SI text. In comparison to the MMC cascade, which was discussed in the previous section, the third cascade element *P*_3_ is more evenly distributed in the cell and concentration gradients are much more shallow (see Figure 2). In the case *γ* = 0, we can derive a expressions for the steady states of the average concentration of signaling components, which are given by

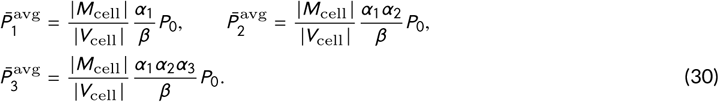

Therefore, the average concentration of the third cascade element 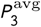 takes the same values in the MMC and PC cascade. The major distinction of both spatial motifs is given by the fact that the concentration differences obtained at the cell membrane and nucleus are larger in the MMC cascade than in the PC cascade. Similarly as in the previous section, we can formulate a system of ordinary differential equations for the average concentrations

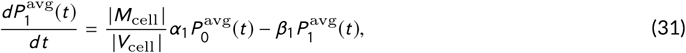

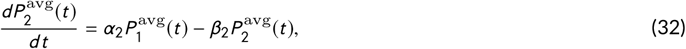

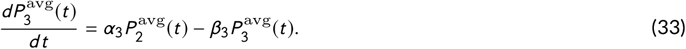

The dependence of absolute concentration levels on the membrane-cytosolic interface is shown in Figure 2 and supplementary Figure S1 for a set of different parameters.

### 3.3 The timing of spatial signaling

The timing of signal transduction in linear signaling cascades for well-stirred homogeneous systems has been analyzed in Heinrich et al. (2002). It was shown that phosphatases have a more pronounced effect than kinases on the rate and duration of signaling, whereas signal amplitude is controlled primarily by kinases. A thorough analysis of linear models assuming a homogeneous distribution of signaling molecules for different kinds of external stimuli has been recently worked out in Beguerisse-Díaz et al. (2016). Here, we want to extend and compare these findings to spatial signal transduction omitting the simplification of homogeneous concentrations. The time-scale analysis for spatial models is more difficult and therefore we used the recently introduced measure of accumulation times Berezhkovskii et al. (2010).

**TABLE 1.** An overview on values and parameters. The numerical values given in the paper all use the units shown in this table.

| entity | value | unit | description |
| --- | --- | --- | --- |
| $D_{\text{mem}}$ | $3.0 \times 10^{-2}$ | $\mu\text{m}^2/\text{s}$ | diffusion coefficient for the membrane-bound species |
| $D_{\text{cyt}}$ | 3.0 | $\mu\text{m}^2/\text{s}$ | diffusion coefficient for the cytosolic species |
| $\alpha_i$ | - | $\text{s}^{-1}$ | phosphorylation rate |
| $\beta_i$ | - | $\text{s}^{-1}$ | phosphatase activity |
| $\gamma$ | - | $\mu\text{m}/\text{s}$ | reaction rate at the membrane- cytosolic interface<br>at the cell membrane |
| $\epsilon$ | - | $\mu\text{m}/\text{s}$ | import rate/permeability at the membrane- cytosolic inter-<br>face at the nucleus |
| $R_{\text{cell}}, R_{\text{nuc}}$ | - | $\mu\text{m}$ | radius of the cell/nucleus |

**TABLE 2.** An overview on values and parameters. The numerical values given in the paper all use the units shown in this table.

The fraction of this steady state level that accumulated at distance *r* and time *t* is expressed as

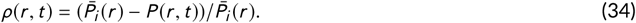

The difference *ρ*(*r*, *t* _1_) - *ρ*(*r*, *t* _2_) can be interpreted as the fraction of the steady state level 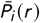 that accumulated in the time interval [*t* _1_, *t* _2_]. In an infinitesimal time interval [*t*, *t* + *d t*] the fraction of accumulated activated signaling molecules at steady state is given by 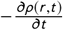. Note, that 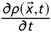 satisfies.

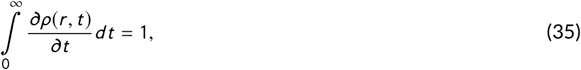

since the initial condition at time *t* = 0 is given by *P*_*n*_ (*r*, 0) ≡ 0. Based on this expression the local accumulation time is defined as Berezhkovskii et al. (2010)

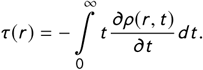

The accumulation time can be derived from the steady state solution even if no closed form of the time-dependent solution is known Berezhkovskii et al. (2010). The timing of the average concentrations given in the system of ordinary differential equations for the MMC cascade (19) - (21) and the PC cascade (31) - (33) are the same and can be analytically expressed as

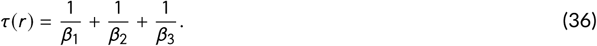

This expression also coincides with signaling times calculated by Heinrich et al. Heinrich et al. (2002). However, for the spatial model the local accumulation times at the membrane and nucleus differ. They are generally faster at the membrane and slower at the nucleus, where the degree of the difference increases with cell size (see Figure 3). Furthermore, also the two spatial motifs show significant differences. For the MMC cascade the accumulation time for the second element *P*_2_ is exactly 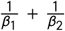 on the membrane, while it is faster for the cytosolic species (compare Figure 3). The accumulation time of *P*_3_ at the nucleus is, as expected, much longer. For a small cell with a Thiele modulus of F < 1, the intracellular concentration is spatially homogeneous and the approximation 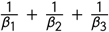 holds, while for signal propagation to the nucleus increases with cell size. An analytical solution of the accumulation times for *P*_3_ for the MMC cascade and the special case of *R*_nuc_ = 0 can be derived Ellery et al. (2013), which is given in the SI Text. The accumulation time for the signal at the nucleus is, as expected longer. For a small cell with F < 1, the intracellular concentration is spatially almost homogeneous and again the approximation 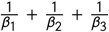 holds. However, for larger cells, the time for signal propagation to the nucleus increases with cell size. For the PC cascade, the increase in accumulation time at the nucleus with cell size is less pronounced than for the mixed-membrane cytosolic cascade.

**FIGURE 3.**
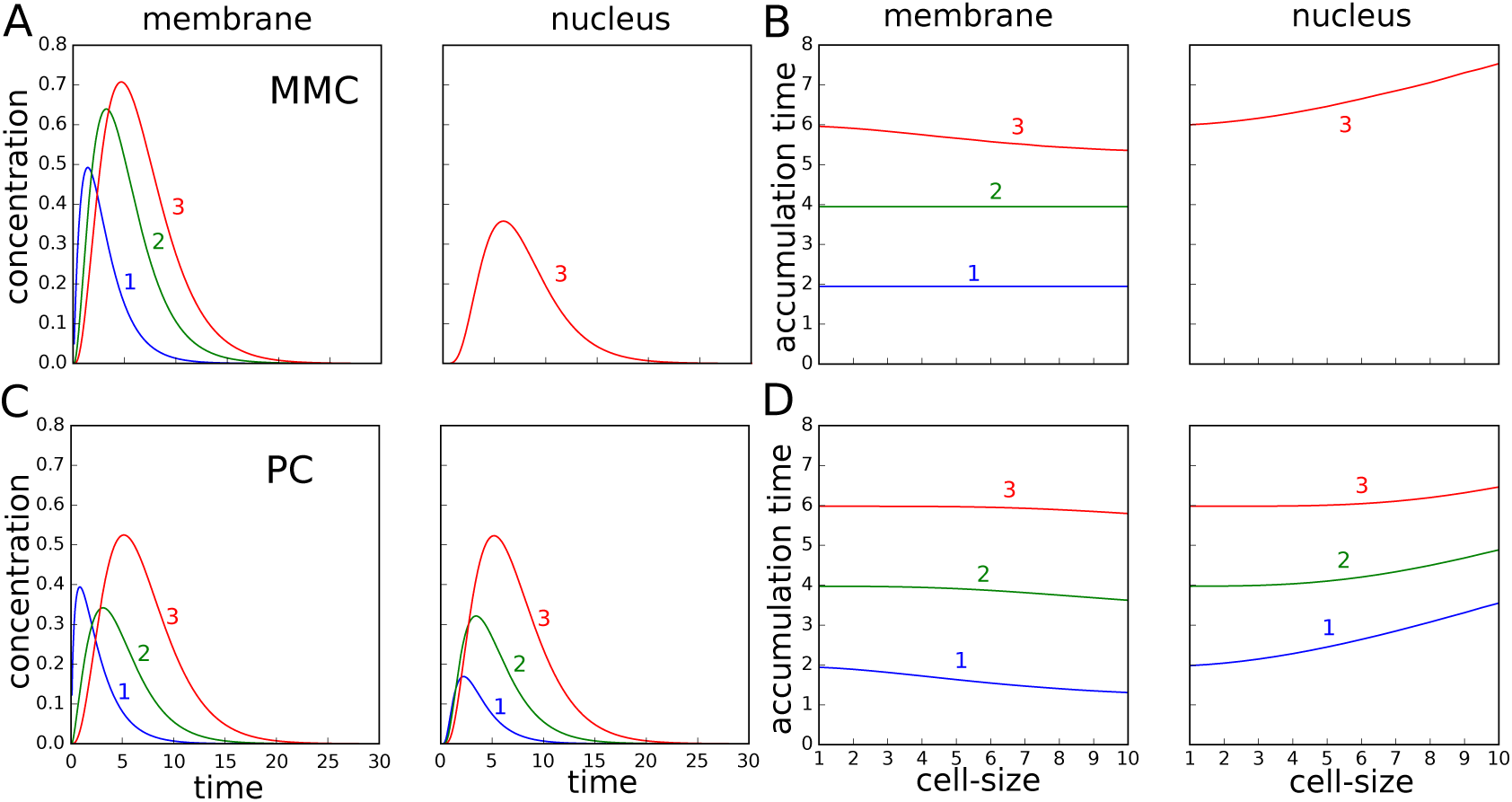
Signaling time at the membrane and nucleus for the mixed membrane-cytosolic (MMC) cascade (**A,B**) and pure cytosolic (PC) cascade (**C,D**) at the membrane and at the nucleus. **A**: Time course of the concentration of *P*_1_, *P*_2_ and *P*_3_ for the mixed membrane-cytosolic cascade. The cascade levels are indicated by the numbers. The signal *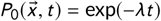* was applied and the parameters used were *R*_cell_ = 6*μm*, *R*_nuc_ = 2*μm*, *λ* = 1, *ɑ*_1_, *ɑ*_2_, *ɑ*_3_ = 1 and *β*_1_, *β*_2_, *β*_3_ = 0.5. **B**: Simulation of the pure cytosolic cascade, but otherwise the same setup and parameters as in (**A**). **C**: Accumulation times for the mixed membrane-cytosolic cascade. In this scenario, a constant signal 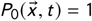 was applied and the cells size varied. The ratio of cellular to nuclear radius was kept at *R*_cell_/*R*_nuc_ = 3. Otherwise the same parameters as in (**A**) and (**B**) were used. **D**: Simulation of the pure cytosolic cascade, but with the same setup and parameters as in (**C**),

While a constant stimulus was applied to calculate the accumulation times, we also tested a decaying signal *P*_0_(*t*) = exp(-*λt*). A comparison of the MMC and PC is shown in Figure 3. Interestingly, the concentration level at the membrane for the PC cascade decreases from the first level *P*_1_ to the second level *P*_2_ and than increases again from the second *P*_2_ to the third *P*_3_ level, while there is an increase from cascade level to cascade level at the nucleus. This phenomenon is caused by the concentration differences from cell membrane to nucleus, which is larger for *P*_1_ than for *P*_2_ in the PC cascade. Note, that the parameters were chosen as 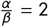), which means a twofold increase for the average concentration levels from one signaling cascade element to the next. Therefore, the spatial system can behave entirely different than the homogeneous system.

For calculation of higher moments of the time scaling and the special case of a cell without nucleus we refer to Ellery et al. (2013). An analysis for time scaling of a linear cascade in one spatial dimension with four elements including higher moments has been carried out in Simpson et al. (2013).

### 3.4 Quantifying the pathway sensitivity with respect to spatially heterogeneous signals

In the following we present a method for analyzing the signal transduction of heterogeneous signals. The signaling cascade can be spatially localized as in the case of directed growth. Examples are *S. cerevisie* Maeder et al. (2007) or *S. pombe* Dudin et al. (2016). Biochemical properties of protein-protein interactions and morphological properties can be tightly connected Peletier et al. (2003). We test the linear signaling cascade with a graded stimulus of the form

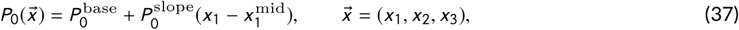

where *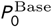* and *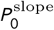* are constants describing the basal signal strength and the slope of the signal, respectively. Here we choose the origin coordinates to be in the center of the cell and, therefore, *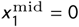*.

The gradient can naturally be defined as the difference of concentration at two points over the euclidean distance of these two points. In the case of the kinase concentrations, the gradient can be computed from *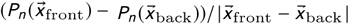*. However, to compare gradients of kinases at different levels, we calculated a normalized gradient in order to eliminate signal amplification of the total concentration

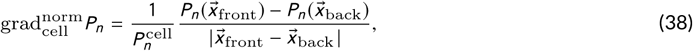

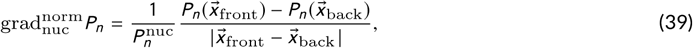

where 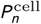 and 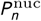 are the average concentrations on the cell membrane and at the nucleus, respectively. For both spatial motifs we performed a parameter study (see Figure 4).

**FIGURE 4.**
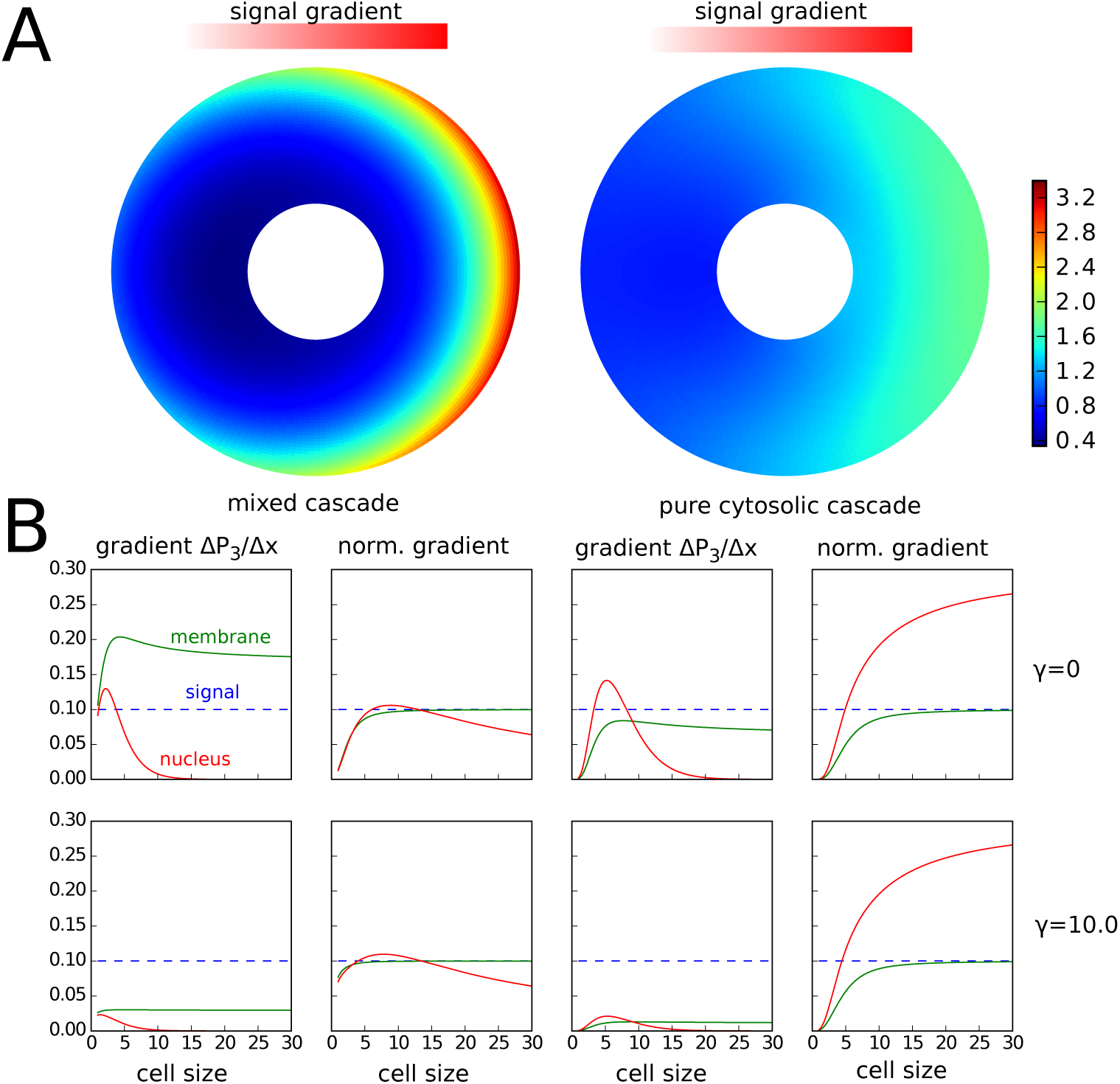
Intracellular concentration profile in response to a linear gradient. The mixed membrane-cytosolic (MMC) cascade with two membrane-bound components and only one cytosolic species exhibits a much stronger response [left] than the the pure cytosolic (PC) cascade, where all three components diffuse freely in the cytosol. Interestingly, for the gradient that is sensed at the cell surface can be higher than the gradient of the signal, which is due to the barrier that the nucleus causes. For the pure cytosolic model theses accumulation effects are most pronounced for the gradient at the nucleus. For both spatial motifs the spatial response to the gradient decreases with higher values of *γ*. The parameters used were *ɑ*_1_ = *ɑ*_2_ = *ɑ*_3_ = 2.0, *β*_1_ = *β*_2_ = *β*_3_ = 1.5, *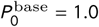* and *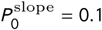*.

Interestingly, the gradient of the third cascade level *P*_3_ first increases and than decreases for both spatial motifs. This effect is much more pronounced at the nucleus than at the membrane. It can be explained by the effect that for a small cells, the concentration is almost homogeneous in the cytosol and concentration differences are balanced by diffusion. However, with increasing cell size the absolute concentration level decreases in the cell and at the nucleus. As a consequence, also the absolute gradient decreases. The normalized gradient at the nucleus for the PC cascade behaves qualitatively similar for the MMC cascade, however the peak for the normalized gradient is obtained for larger cell sizes. For the PC cascade the normalized gradient is increasing with cell size in the observed range of cell sizes.

The findings can also be generalized to higher order spatial heterogeneities, meaning heterogeneities with multiple maxima and minima (see SI Text). The signal can be decomposed using spherical harmonics

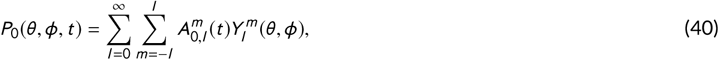

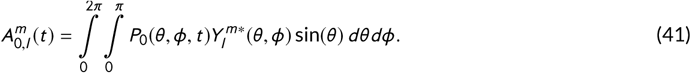

In this decomposition the amplitudes of higher order, where the order is denoted by *l*, are generally more strongly damped than gradients or spatial heterogeneities of lower order (see SI Text). In this manner, the results shown here can be extended to complex spatial signals on the cell surface.

Furthermore, we tested the influence of asymmetries in cell shape and organelle position. The first asymmetry is a cell with a nucleus moved away from the cell center (see Figure 5). The second asymmetry is a cell with a protrusion, as occurs for instance in mating yeast Diener et al. (2014). Both cellular asymmetries induce an intracellular gradient along the cell membrane as well as the nuclear membrane even if a spatially homogeneous gradient is applied on the cell surface (see Figure 5**B**). For a cell with the nucleus moved away from the center a gradient is induced, since the nucleus acts like a dam against the diffusive flux from the membrane into the cytosol (see Figure 5**B**). For the cell with a protrusion the concentration of *P*_3_ is higher in the protrusion than in the opposite distal end, which is the spherical part of the cell. This effect emerges due a higher local surface to volume ratio in the protrusion region. Therefore, a larger portion of cytosolic signaling molecules, which freely diffuse in the cytosol, is phosphorylated in the protrusion part leading to a gradient from the protrusion tip to the opposite distal end of the cell. The influence of cellular asymmetries has also been investigated in Giese et al. (2015) for gradients of the small Rho-GTPase Cdc42 during cell polarization. However, this system reacts in the opposite way since the flux of molecules is directed from the cytosol onto the membrane and one observes a gradient of from the distal end to the protrusion. For a cell with an organelle moved away from the cell center, the polarization system results in a lower concentration in the vicinity of the organelle. These effects occur due to the different architectures of both systems. In the PC and MMC signaling cascades, we have signal transduction from the membrane to the nucleus and, therefore, a diffusive flux of activated signaling molecules from the membrane into the cytosol, while in the polarization system the flux of signaling molecules is directed from the cytosol onto the membrane, which is the opposite direction. Therefore, both system respond differently to cellular asymmetries with respect to gradient formation. This interplay of both systems is especially interesting, since in many organisms a polarization system is interacting with a MAPK cascade Thomson et al. (2011); Ventura et al. (2014). To complete our investigations, we varied the slope of the signal and investigated the sensitivity of both cell shapes towards different slopes of signals and compared the response with a symmetric cell (see Figure 5**C**).

**FIGURE 5.**
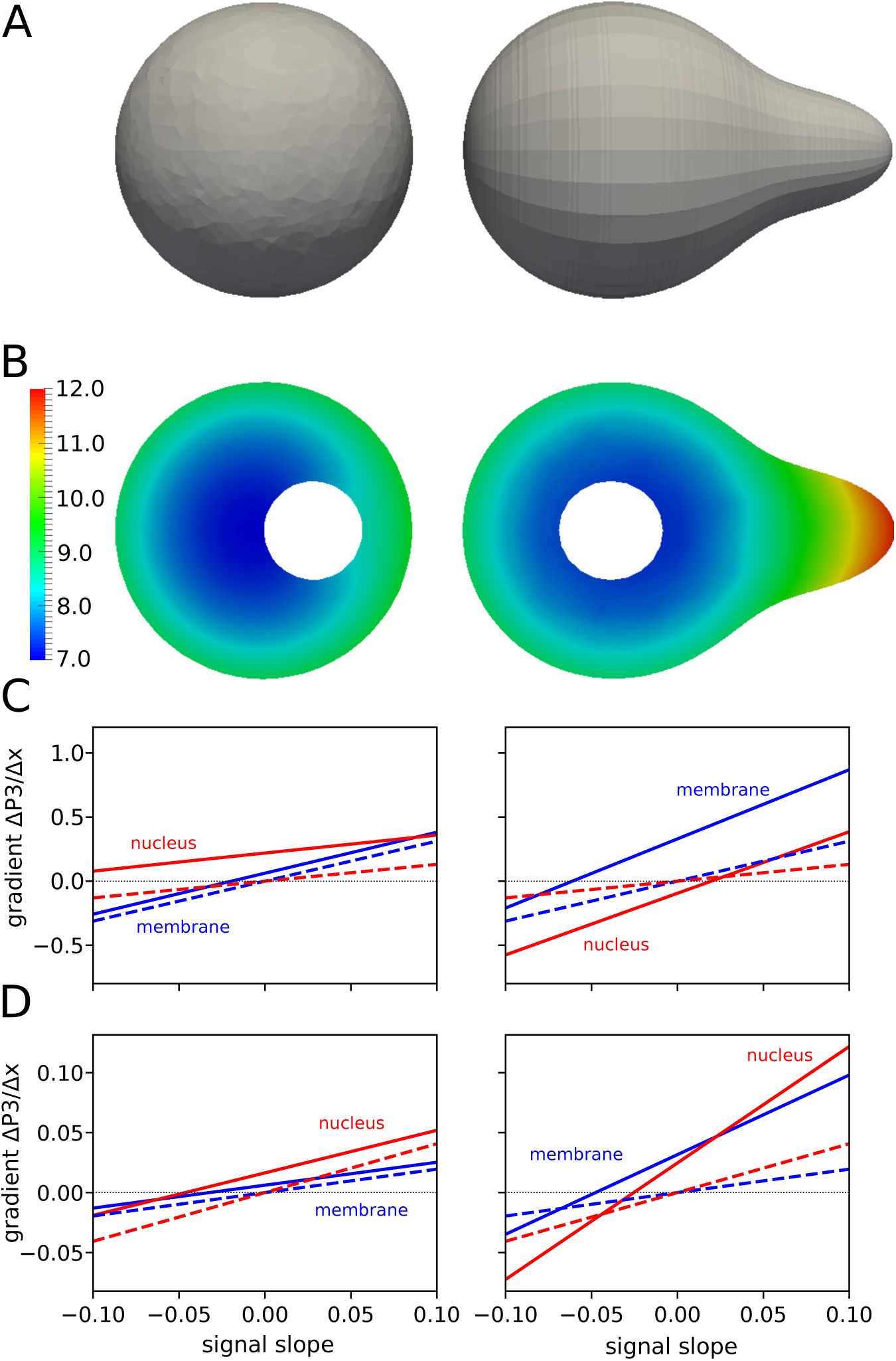
Response of the two cascade systems (MMC and PC) for different cellular asymmetries to a linear gradients with varying slope *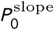*. **A** and **B**: The two investigated cell shapes. A spherical cell with *R*_cell_ = 3 *μm*, *R*_nuc_ = 1 *μm* and a nucleus shifted to the side as well as a cell with a protrusion. **B**: Profile of *P*_3_ after stimulation with a spatially homogeneous signal, meaning *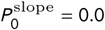*. Due to the cellular asymmetry in organelle position (left) and asymmetric cell shape (right) a gradient along the cell membrane and nuclear membrane is generated, which would not be generated for a symmetric cell. **C**: Response to different values of *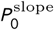* for the MMC cascade. The response of a symmetric cell with *R*_cell_ = 3 *μm* and *R*_nuc_ = 1 *μm* is indicated with dashed lines for comparison. The gradient of *P*_3_ in case of the cellular asymmetry is altered (solid lines). **D**: Same setup for the PC cascade. The parameters used were *ɑ*_1_ = *ɑ*_2_ = *ɑ*_3_ = 1.0, *β*_1_ = *β*_2_ = *β*_3_ = 0.5, *γ* = 0.0 and *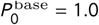*.

## 4 DISCUSSION

Stimulated by the progress in cell imaging and the increasing need to understand intracellular dynamics, we investigated and discussed a general approach of modeling cellular signal transduction in time and space. We showed that modeling of the membrane-cytosolic interface is crucial as well as the ratio of membrane area and cytosolic volume, which are both spatial properties. The results imply strong cell size dependence of signal transduction within cells. Widely used time-dependent models of ordinary differential equations can naturally be extended into space by using bulk-surface differential equations. Applying this extension to a class of linear signal transduction models, we compared the assumption of a well mixed cell with two different spatial signal transduction motifs. We derived and discuss criteria that can be used to test the well-mixed assumption and show that kinetics that connect membrane-bound species with cytosolic species naturally cause size dependence. The results are therefore of general importance for kinetic models of signal transduction.

Our findings have relevant biological implications. Since the signals transduced by linear signaling cascades from the cell membrane to the nucleus decrease exponentially on a length scale of a few microns, our theoretical findings suggest a strong cell size dependence in response to extracellular stimuli. Interesting studies of the response in cell populations often lack the response behavior attributed to cell size and morphology. Examples range from the switch-like behavior in populations of oocytes Ferrell and Machleder (1998) to the pheromone response in yeast cells Conlon et al. (2016); Banderas et al. (2016). Therefore, single cell data where the cell size is assigned to the measurements is needed for a faithful quantitative investigation of the pathway, to disentangle biochemical properties of protein-protein interactions to morphological properties such as size and shape of whole cells.

In non-linear systems, the differences that we observed in the linear signaling cascade models are likely to be amplified. Non-linear kinetics can amplify gradient formation, which leads to even stronger intracellular concentration differences Wartlick et al. (2009). This also holds for absolute concentration levels that can behave in a switch-like manner depending on the kinetics Kholodenko (2000); Ferrell and Machleder (1998). The same holds for the time scales of signaling. Higher order kinetics can amplify the accumulation time differences in different cellular locations Gordon et al. (2011).

The model can be extended to more complex spatial heterogeneities for example by using the Laplace series as suggested in this paper. With localized signals arising from membrane structures like lipid rafts, septins, co-localization due to protein-protein interactions can be represented. Since these are often precursors for cell shape and organelle structures the interplay with cell shape needs to be addressed by future research. The intrinsic geometry dependence of bulk-surface signaling systems has recently been shown for ellipsoidal cell shapes in the MinE-MinD system Halatek and Frey (2012); Thalmeier et al. (2016); Wu et al. (2016), but also in the yeast system Orlandini et al. (2013); Giese et al. (2015); Chen et al. (2016). Here, not only the global but also the local surface to volume ratio in cell protrusion plays an important role. Recent developments of mathematical methods such as the finite element method for bulk-surface equations Elliott and Ranner (2012); Eigel and Müller (2017) as well as stability analysis techniques Rubinstein et al. (2012); Edelstein-Keshet et al. (2013); Rätz and Röger (2014); Giese et al. (2015); Garcke et al. (2016); Madzvamuse et al. (2016) are expected to provide further insight in the behavior of biological systems.

## METHODS

We used the finite-element software FEniCS Alnæs et al. (2015); Logg et al. (2012) to generate the corresponding meshes and to solve the arising partial differential equations in the Python programming language. The non-linear equations were solved using a fixed-point scheme Milicic et al..

## AUTHOR CONTRIBUTIONS

Conceptualization, EK and WG; Methodology, EK, WG, GM and AS; Software, WG and GM; Writing – Original Draft, EK, WG and GM; Writing - Review & Editing, EK, WG, GM and AS; Supervision, EK, WG and AS; Funding Acquisition, EK and AS.

## Abbreviations

*MAPK*: mitogen-activated protein kinase
*PC*: pure cytosolic
*MMC*: mixed membrane-cytosolic
*PDE*: partial-differential equation

